# The EXS domain of rice PHOSPHATE1 (OsPHO1;2) reduces the low phosphate response through jasmonate signaling

**DOI:** 10.1101/2025.11.07.687324

**Authors:** Balaji Mani, Kanika Maurya, Lokesh Verma, Priya Gupta, Pawandeep Singh Kohli, Gagan Gupta, Aime Jaskolowski, Yves Poirier, Jitender Giri

## Abstract

Phosphorus (P) is a vital macronutrient essential for plant growth, and its deficiency significantly hampers agricultural productivity. The PHOSPHATE 1 (PHO1) protein family, characterized by an N-terminal SPX domain, four transmembrane (4TM) domains, and a C-terminal EXS domain, is pivotal in transporting phosphate (Pi) from roots to shoots. Rice, PHO1;2 plays a crucial role in the Pi export process, and defects in this gene cause severe growth retardation and Pi deficiency symptoms even when external Pi levels are adequate. This study examined the roles of the EXS domain and the combined 4TM+EXS domains of OsPHO1;2 in supporting plant growth responses, independent of Pi transport activity, as well as their influence on hormone signaling and gene regulation. Using CRISPR/Cas9, rice lines expressing specific OsPHO1;2 domains (EXS or 4TM+EXS) were created by targeted deletion of particular domain-coding regions. Phenotypic analysis under Pi-sufficient and deficient conditions, as well as phosphate profiling, revealed that EXS lines exhibited notably better growth than loss-of-function mutants, *ospho1;2,* during early development, despite having similar shoot Pi levels to the null mutant. These lines showed lower levels of defense hormones (jasmonic acid) than *ospho1;2* but were comparable to those in the wild type. Conversely, 4TM+EXS lines exhibited growth patterns similar to *ospho1;2* mutants. RNA sequencing indicated that the phosphate starvation response (PSR) and defense pathways were less pronounced in the EXS lines compared to *ospho1;2* mutants. However, both EXS and 4TM+EXS lines showed seed development defects and reduced total phosphorus content in seeds, mirroring the *ospho1;2* phenotype. Heterozygous plants carrying one functional *OsPHO1;2* allele displayed normal growth and seed development, indicating the mutation’s recessive nature. The findings suggest that the EXS domain of OsPHO1;2 can promote plant growth independently of Pi transport by decreasing PSR and modulating defense hormone pathways. This further suggests a signaling role for PHO1 domains beyond direct Pi translocation. Overall, these results enhance our understanding of Pi homeostasis and may help form strategies for breeding P-efficient crops.

## Introduction

Phosphorus (P) is a macronutrient for vital biological processes, like energy metabolism, signal transduction, and the structural components of nucleic acids and phospholipids(Kopriva & Chu, 2018). The primary form of P available to plants is inorganic orthophosphate (Pi), which must be efficiently imported from the soil solution and distributed throughout plant tissues to support optimal growth and development (Yang et al., 2024). Plants have developed diverse adaptations to survive under Pi starvation (Plaxton & Tran, 2011). One such adaptation involves the evolution of complex Pi transporters, broadly classified into families such as phosphate transporters (PHTs), PHOSPHATE1 (PHO1), and SULTR-like phosphorus distribution transporter (SPDT), which facilitate Pi acquisition and distribution to growing tissues/organs (Yang et al., 2024). Understanding how plants manage Pi transport and exploring strategies to improve Pi acquisition and utilization are crucial for increasing crop productivity. The PHO1 family is a unique group of Pi exporters that are essential for long-distance Pi transport from roots to shoots. PHO1 proteins are characterized by their distinctive domain structure: an N-terminal SPX (SYG1/Pho81/XPR1) domain, four transmembrane (4TM) domains, and a C-terminal EXS (ERD1/XPR1/SYG1) domain (Wang et al., 2004; Wege et al., 2016). This multi-domain structure allows PHO1 to function both as a Pi transporter and as a component of phosphate signaling networks (Poirier et al., 1991; Wild et al., 2016).

The SPX domain acts as an intracellular sensor of the plant’s Pi status by interacting with inositol polyphosphates (InsPs) to regulate the expression of phosphate starvation response (PSR) genes (Wild et al., 2016; Dong et al., 2019; Ried et al., 2021; Wang et al., 2021). The EXS domain, located within a C-terminal hydrophobic region, contains key structural elements necessary for Pi export activity and proper subcellular localization of Arabidopsis PHO1 protein (AtPHO1) (Wege et al., 2016). The AtPHO1 is localized in the Golgi/Trans-Golgi network (Arpat et al., 2012). The truncated AtPHO1, lacking the EXS domain, causes the protein to remain in the endoplasmic reticulum (Wege et al., 2016). However, transient expression of the AtPHO1 EXS domain in *Nicotiana benthamiana* directs it to the Golgi/Trans-Golgi network, similar to AtPHO1 (Wege et al., 2016). Additionally, the EXS domain is essential for trafficking PHO1 to and from the plasma membrane, thus helping regulate Pi homeostasis (Vetal & Poirier, 2023). Mutants of AtPHO1 show impaired Pi unloading into root xylem tissues, leading to Pi-deficient shoots and reduced plant growth (Poirier et al., 1991; Wang et al., 2004; Secco et al., 2010). These phenotypes are accompanied by increased expression of phosphate starvation-induced (PSI) genes and accumulation of phytohormones like jasmonic acid (JA) (Khan et al., 2016) and abscisic acid (ABA) in shoots (Jaskolowski & Poirier, 2024). AtPHO1 is also involved in the export of Pi from the maternal chalazal seed coat to the filial embryo (Vogiatzaki et al., 2017).

In rice (*Oryza sativa*), the PHO1 family includes three members (*OsPHO1;1*, *OsPHO1;2*, and *OsPHO1;3*), with *OsPHO1;2* playing the main role in root-to-shoot Pi translocation (Secco et al., 2010). *OsPHO1;2* is mainly expressed in roots, and its loss-of-function causes severe growth defects that resemble Pi deficiency symptoms even under Pi-sufficient conditions due to impaired Pi export (Secco et al., 2010; Ma et al., 2021). Ma *et al*. (2021) further demonstrated that the *ospho1;2* mutation has a significant impact on seed development, yield, and plant growth. Conversely, overexpressing *OsPHO1;2* boosts Pi transport, increases panicle number, overall yield, phosphate utilization efficiency (PUE), and photosynthesis performance, indicating its potential to improve plant Pi status and overall growth under different Pi conditions (Ma et al., 2021, 2024). In the development of rice seeds, OsPHO1;2 is expressed in the maternal caryopsis coat and is involved in the export of Pi to the seed filial tissues (Ko et al., 2024). Recent research has revealed that removing the repressor site (*W-box*) in the *OsPHO1;2* promoter greatly enhances *OsPHO1;2* expression, improves Pi export from roots to shoots, and increases yield (Maurya *et al*. 2025).

The interaction between phytohormones and Pi signaling influences plant growth and development (Pandey et al., 2021; Mehra et al., 2022). Pi deficiency triggers defense signaling and results in the accumulation of jasmonic acid (JA), salicylic acid (SA), and abscisic acid (ABA) (Khan et al., 2016; Zhang et al., 2022a; Wu et al., 2022; Jaskolowski & Poirier, 2024). The *atpho1* mutants grown under Pi-sufficient conditions exhibit higher JA levels, show poor growth, but display resistance to *Spodoptera* infection (Khan et al., 2016). Conversely, the Pi deficiency of the *atpho1* mutants makes them more vulnerable to *Botrytis cinerea* infection (Jaskolowski & Poirier, 2024). This is mainly due to higher ABA levels in *atpho1*, which enhance susceptibility to *Botrytis cinerea* by promoting spore adhesion and germination and reducing callose deposition. This suggests that both JA and ABA are involved in low Pi-mediated signaling in plants.

In Arabidopsis, the EXS domain alone can restore shoot growth in *atpho1* mutants while maintaining low shoot Pi levels, indicating that it plays a role in root-to-shoot signaling that distinguishes growth defects from Pi deficiency symptoms (Wege et al., 2016). A similar trend of low shoot Pi with enhanced growth is also seen in AtPHO1 gene-silenced lines (Rouached et al., 2011), suggesting that reduced AtPHO1 expression or the expression of EXS alone can separate growth defects from Pi deficiency symptoms. However, the molecular mechanisms underlying the uncoupling of Pi deficiency symptoms from growth defects remain unclear, especially in cereal crops like rice, where Pi efficiency is crucial for global food security. Understanding how different PHO1 domains influence plant growth, Pi homeostasis, and defense hormone signaling could provide valuable insights for developing crops with enhanced phosphorus use efficiency. Additionally, studying the evolutionary conservation of EXS domain functions across plant species would deepen our understanding of fundamental Pi signaling pathways and their potential in crop improvement.

In the current study, we aimed to edit rice PHO1;2 so that it expresses only the EXS domain under the *OsPHO1;2* promoter, which is expected to result in better growth under limited Pi conditions. By creating genome-edited rice lines that express specific *OsPHO1;2* domains, we uncovered a role for *OsPHO1;2* in Pi signaling via the EXS domain. Our findings improve the understanding of PHO1 domain-specific functions and their potential for developing Pi-efficient crops for sustainable agriculture.

## Materials and methods

### Plant materials and growth conditions

Rice (*Oryza sativa ssp. Japonica*) was used in all experiments and employed for generating CRISPR/Cas9 rice lines. For phenotypic, molecular, and physiological analyses, dehusked rice seeds were surface-sterilized with a 0.1% HgCl2 solution and germinated on half-strength Murashige and Skoog (MS) medium for five days. After germination, the seedlings were transferred to phosphate-deficient media (50 μM NaH_2_PO_4_) and phosphate-sufficient media (320 μM NaH_2_PO_4_), as described by (Maurya et al., 2025). All rice lines were grown in a controlled growth chamber with a day/night temperature of 30 °C/28 °C, a photoperiod of 16 hours of light and 8 hours of darkness (400-450 μmol photons m−2 s−1), and a relative humidity of ∼70%.

### Generation of *OsPHO1;2* EXS lines using CRISPR/Cas9 ribonucleoprotein (RNP) complexes

The rice lines expressing the EXS domain were created by removing the SPX and 4TM domains (EXS lines) using RNP complexes. Two guide RNAs (gRNAs) targeting the region flanking the SPX and 4TM domains coding sequence of *OsPHO1;2* were designed with the CRISPR-GE toolkit (http://skl.scau.edu.cn/home/). In vitro transcripts of gRNA were synthesized using the EnGen sgRNA Synthesis Kit (NEB, USA), following the manufacturer’s instructions. After DNase treatment, the transcribed gRNAs were purified with NucAway spin columns (Invitrogen, USA).

For Cas9 protein expression, the Cas9 coding sequence, along with SV40 and nucleoplasmin nuclear localization signals, was amplified from the pRGEB32 vector and cloned into the pET28a vector (**Fig.S1A**). The Cas9 protein was expressed in the *E. coli* Rosetta strain at 18°C, followed by purification using cOmplete™ His-Tag Purification Resin (Roche, USA), and concentrated with an Amicon® Ultra Centrifugal Filter (Millipore, USA). The purity and concentration of the Cas9 protein were determined by SDS-PAGE (**Fig.S1B**) and the Bradford protein assay, respectively. All primer sequences used are listed in **Table S1**.

The formation of the RNP complex and particle bombardment were carried out according to the protocol described by Liang et al. (2017), with minor modifications. For each shot, the RNP complex was prepared by mixing equal molar amounts of gRNAs (2 µg) and Cas9 protein (2 µg), followed by incubation at room temperature for 15 minutes. The RNPs were combined with gold particles (0.6 mm) (Bio-Rad, USA) and kept on ice for 10 minutes. The RNP-coated gold particles were pelleted at 6,000 g for 1 minute, then resuspended in 20 µL of sterile water and loaded onto a macrocarrier to air-dry for 1 to 2 hours. 10-day-old rice calli derived from the scutellum were bombarded using a PDS-1000/He Gun (Bio-Rad, USA) set at a rupture pressure of 1100 PSI. After bombardment, the calli were transferred to ½ MS media and incubated at 28°C for 5 days, after which plant regeneration proceeded as described by Maurya *et al*. (2025).

### Vector construction and plant transformation

To create *ospho1;2* knock-out (KO) lines, a gRNA targeting the first exon of *ospho1;2* was identified using the CRISPR-GE toolkit (http://skl.scau.edu.cn/home/). This gRNA was mobilized into the pRGEB32 vector, and transgenic rice lines were produced through *Agrobacterium tumefaciens-*mediated transformation, as mentioned (Maurya *et al.,* 2025). The sequences of all the primers used are listed in **Table S1.**

The *OsPHO1;2* line, which expresses four TM domains plus the EXS domain (4TM+EXS), was created by deleting the SPX domain coding sequence. The two gRNAs flanking the SPX domain of *OsPHO1;2* were identified, and a tRNA-gRNA-tRNA-gRNA (PTG) complex was assembled (Xie et al., 2015). This PTG complex was then inserted into the pRGEB32 vector and used to develop 4TM+EXS lines via *Agrobacterium tumefaciens-*mediated transformation, as described (Maurya *et al.,* 2025). The mutants were confirmed through genomic DNA PCR and Sanger sequencing. All primer sequences are listed in **Table S1**.

### Quantitative real-time PCR analysis

Total RNA isolation, cDNA synthesis, and quantitative real-time PCR (qRT-PCR) were conducted according to the methods described by (Mani et al., 2024). The rice eEF1a (*eukaryotic translation elongation factor 1a*) gene was used as the internal control. The nucleotide sequences of the primers used for gene expression analyses are listed in **Table S1.**

### Transcriptome sequencing and analysis

Five-day-old seedlings of WT, *ospho1;2*, and EXS-1 lines were transferred to Pi-sufficient media (320 µM) and grown for 14 days. Total RNA was extracted from the shoot tissues using the Direct-zol RNA kit (Zymo Research, USA) and quantified with NanoDrop One (Thermo Fisher Scientific, USA). RNA integrity was assessed with the Plant RNA Nano Assay Kit on the Agilent Bioanalyzer 2100 system. Samples with an RNA integrity number (RIN) above 7.0 were used for library preparation using the KAPA RNA HyperPrep Kit (Roche, USA). Three biological replicates for WT, *ospho1;2*, and EXS-1 lines were sequenced on the Illumina NovaSeq 6000 platform with 151 bp paired-end reads. High-quality clean reads were obtained from raw reads using FastQC (v0.11.9) and fastp (v0.23.2) (Chen et al., 2018). The trimmed paired-end reads were aligned to the reference genome (*Oryza sativa - Nipponbare* - Reference IRGSP-1.0) using Hisat2 (v2.1.0) (Kim et al., 2019). Gene expression was quantified, and differentially expressed genes were identified using FeatureCounts (v1.34.0). DESeq2 (v1.44.0) was employed to identify differentially expressed genes (DEGs) between samples (Love et al., 2014). Genes with a log2 (base 2) value ≥ 1 were considered upregulated, while those with a log2 (base 2) value ≤-1 were considered downregulated, with a false discovery rate (FDR) of < 0.05. GO functional enrichment analysis for GO terms with FDR < 0.05 was performed using ShinyGO. The raw transcriptome data were uploaded to IBDC (https://ibdc.dbtindia.gov.in/ accession number: INRP000478; PRJEB100756).

### Quantification of defense phytohormones

Five-day-old seedlings of WT, *ospho1;2*, and EXS-1 lines were transferred to Pi-sufficient media (320 µM) and allowed to grow for 14 days. The 20 mg of lyophilized shoot samples were used to quantify phytohormones, containing 40 ng/mL of D6-ABA and 8 ng/mL of JA-isoleucine conjugate as internal standards. The samples were shaken for 30 minutes at 4 °C and then centrifuged at 13,000 g for 20 minutes at 4 °C. The supernatant was collected, and the samples were re-extracted with 500 µL of methanol. After centrifugation, the supernatants were combined and evaporated in a SpeedVac at ambient temperature (25 °C) before being re-dissolved in 500 µL of methanol. The samples were analyzed using an Exion LC (Sciex®) UHPLC system, following the methods outlined by (Pandey et al., 2025). Phytohormones were quantified relative to their corresponding internal standards.

### Estimation of soluble Pi and Total P content

The Pi content was determined with the molybdate assay, as outlined by Ames (1966). The total phosphorus concentration in the plant samples was measured using the vanadate–molybdate method (Hanson, 1950).

## Statistical analysis

The data from all qRT-PCR experiments, morphometric, and physiological analyses were analyzed using one-way analysis of variance (ANOVA), followed by Tukey’s honestly significant difference (HSD) test to assess the statistical differences among the various groups. The statistical analysis was conducted using GraphPad Prism 8 software.

## Results

We hypothesize that plants expressing only the EXS domain under the *OsPHO1;2* native promoter would grow well despite having reduced shoot Pi levels. To test this, we made rice *pho1;2* mutants and lines expressing different domains of OsPHO1;2.

### Generating rice lines expressing only the EXS domain under the *OsPHO1;2* native promoter (EXS lines)

Gene architecture analysis revealed that *OsPHO1;2* consists of 15 exons separated by 14 introns, along with short 5’ and 3’ UTRs. To create lines expressing only the EXS domain of OsPHO1;2 (thereafter named EXS lines), we intended to remove the region between the first and eleventh introns, which encode the SPX and four TM domains (**Fig. 1A**), deleting approximately 3.5 kb from the *OsPHO1;2* locus (**Fig. 1A**). In these lines, the first exon, which partially encodes the SPX domain, remains in frame with the exons that encode the EXS domain, thus producing a truncated protein. After screening over 100 lines, we got two lines with the desired EXS genotypes, named EXS-1 and EXS-2 (**Fig. 1B**). Semi-quantitative RT-PCR verified the presence of a truncated transcript of about 1 kb in the EXS lines, while RT-qPCR confirmed domain-specific expression of *OsPHO1;2* in these lines (**Fig. 1C, D**). We obtained a precise deletion of 3461 base pairs (bp) encompassing SPX and 4TM in both EXS lines, leaving only EXS domain (**Fig. 1E**).

**Fig. 1.**
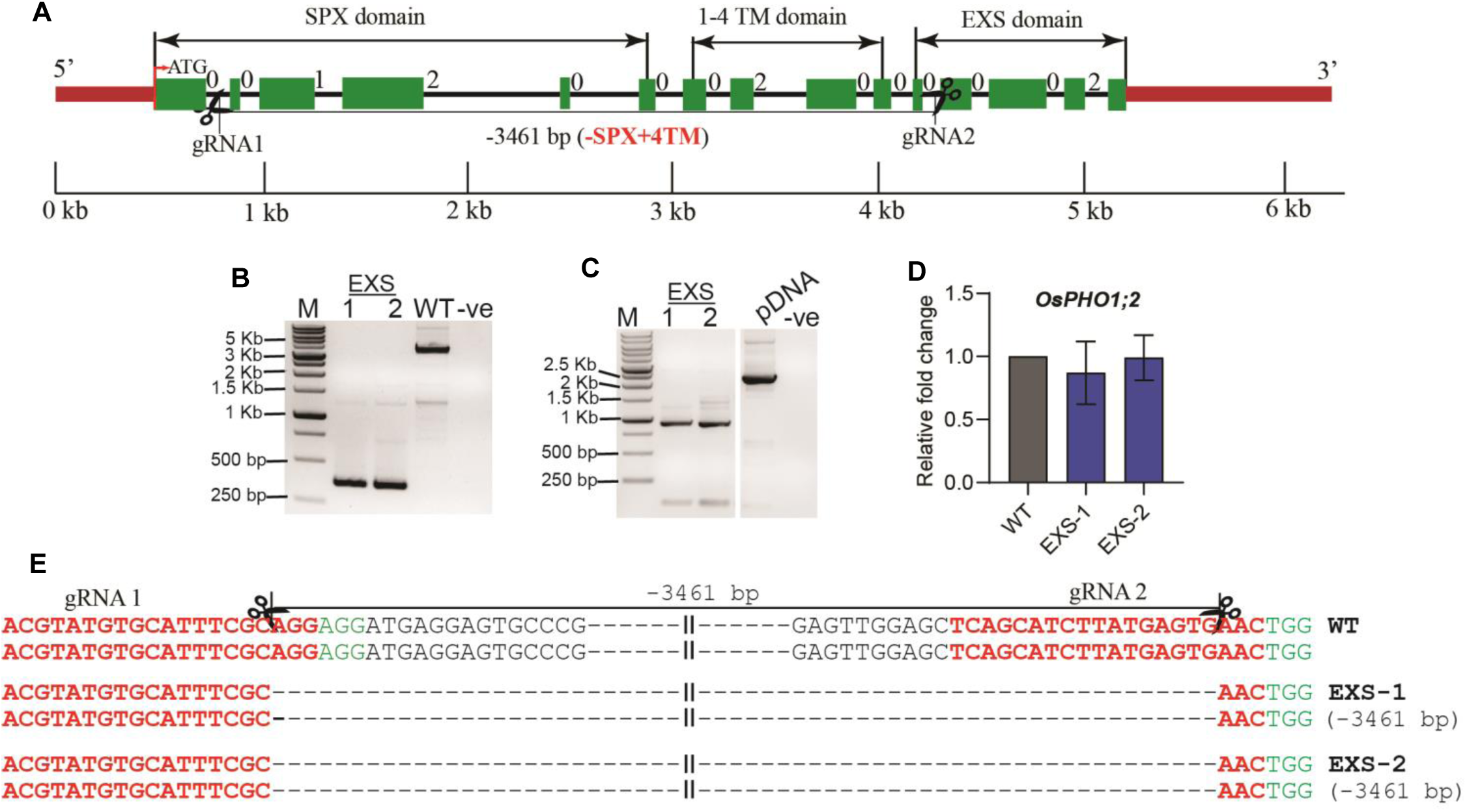
Rice lines expressing only the EXS domain under the OsPHO1;2 native promoter. **(A)** Schematic representation of the intron-exon composition of OsPHO1;2 and gRNA binding positions to the target site for Cas9-mediated targeted gene region deletion. Exons, introns, and UTRs of OsPHO1;2 are represented by green boxes, black and red lines, respectively. Intron phases are defined as 0, 1, and 2. The exons encoding for the SPX domain, transmembrane (TM) domain, and EXS domain are indicated by arrow marks. The specific gRNA/Cas9-mediated cleavage site of the target genes is indicated by scissors, and the expected gene region deletion is depicted below in base pairs (bp). (B) Genotyping of progenies from T2 generation EXS plants and WT by genomic DNA-PCR. M-DNA ladder; bp-base pairs,-ve - PCR negative. (C) Semi-quantitative RT-PCR detection of OsPHO1;2 transcripts in root tissues of WT and EXS genome-edited lines. (D) RT-qPCR analysis of transcript levels of OsPHO1;2 in root tissues of EXS genome-edited lines. The elongation factor 1-_ll_ (EF1_ll_) was used as an internal control for normalizing Ct values. The data is based on three independent biological replicates’ standard error mean values. (E) Sanger-sequencing-based detection of OsPHO1;2 deletions in EXS lines. The 20 bp gRNA targeting sequence and Protospacer adjacent motif (PAM) sequence are shown in red and green colored highlighted nucleotides, respectively. A vertical arrowhead indicates the expected cleavage site, and the expected deletion of the target region is shown on the top. The genotypes and observed target deletion are shown on the left side of the figure. WT represents nucleotide sequence identical to the OsPHO1;2.

To compare the phenotypes of the EXS lines, we created *ospho1;2* knockout (KO) lines by targeting the first exon of *OsPHO1;2* (**Fig. S2A**). DNA sequencing revealed a 7-bp deletion in the *ospho1;2-11* line and a 2-bp deletion in the *ospho1;2-14* line, both resulting in *OsPHO1;2* knockouts (**Fig. S2B, C**). RT-qPCR analysis showed a 70% reduction in *OsPHO1;2* transcript levels in both lines compared to the WT (**Fig. S2D**). In agreement with earlier reports (Secco et al., 2010; Ma et al., 2021), both *ospho1;2* KO lines had poor growth, lower shoot Pi, and higher root Pi levels compared to the WT (**Fig. S2E-K**). Since both KO lines, *ospho1;2-11* and *ospho1;2-14*, exhibited similar behaviors, we selected *ospho1;2-14* for further experiments.

### EXS lines exhibit growth similar to wild-type despite highly reduced shoot P levels

To study the performance of EXS lines, we grew these lines and WT under Pi-sufficient conditions (320 µM NaH_2_PO_4_) for 7 days. The EXS lines had significantly higher shoot length, root length, and better fresh weight (FW) compared to *ospho1;2,* which was severely dwarfed. Importantly, all growth parameters of the EXS lines were comparable to WT (**Fig. S3A-D**) at the young seedling stage, thus supporting our hypothesis. Remarkably, EXS lines exhibited shoot Pi levels similar to *ospho1;2*, but significantly lower than WT, while root Pi levels were higher than WT but comparable to *ospho1;2* (**Fig. S3E, F)**. We further examined the growth of the EXS lines in Pi-deficient conditions (50 µM NaH_2_PO_4_) for 7 days. The EXS lines continued to perform better in growth and biomass accumulation than *ospho1;2* lines (**Fig. S4A-D**) despite highly reduced Pi levels. Their shoot and root Pi levels were similar to *ospho1;2* but differed from WT (**Fig. S4E, F**). Additionally, we assessed the phenotypes of these lines after 14 days under Pi-sufficient conditions. The EXS lines showed a 15% increase in shoot length, a 22% increase in shoot FW, and a 25% increase in shoot dry weight (DW) compared to *ospho1;2*, and were similar to WT (**Fig. 2B-F**). Both *ospho1;2* and EXS lines had significantly lower Pi levels in shoots and higher in roots compared to WT (**Fig. 2g, H**). RT-qPCR analysis revealed higher expression of PSI genes, such as *OsIPS1*, *OsIPS2*, *OsMGD3*, and *OsGDPD5*, in shoot tissues of *ospho1;2* mutants compared to WT. Interestingly, these PSI genes were not upregulated in the EXS lines relative to WT (**Fig. 2I-L**). Furthermore, when grown under Pi-deficient conditions for 14 days, the EXS lines demonstrated better overall growth in shoot length, FW, and DW than *ospho1;2* (**Fig. 3A-G**). Notably, even under Pi deficiency, PSI gene expression was not higher in the EXS lines compared to WT (**Fig. 3H-K**). These findings suggest that, despite accumulating ∼50% less shoot Pi than WT, expression of the EXS domain yields growth like WT and better than *ospho1;2.* This improved growth performance may be attributed to better metabolic adaptations or resource allocation, allowing EXS domain-expressing plants to compensate for lower Pi levels and achieve growth comparable to WT plants.

**Fig. 2.**
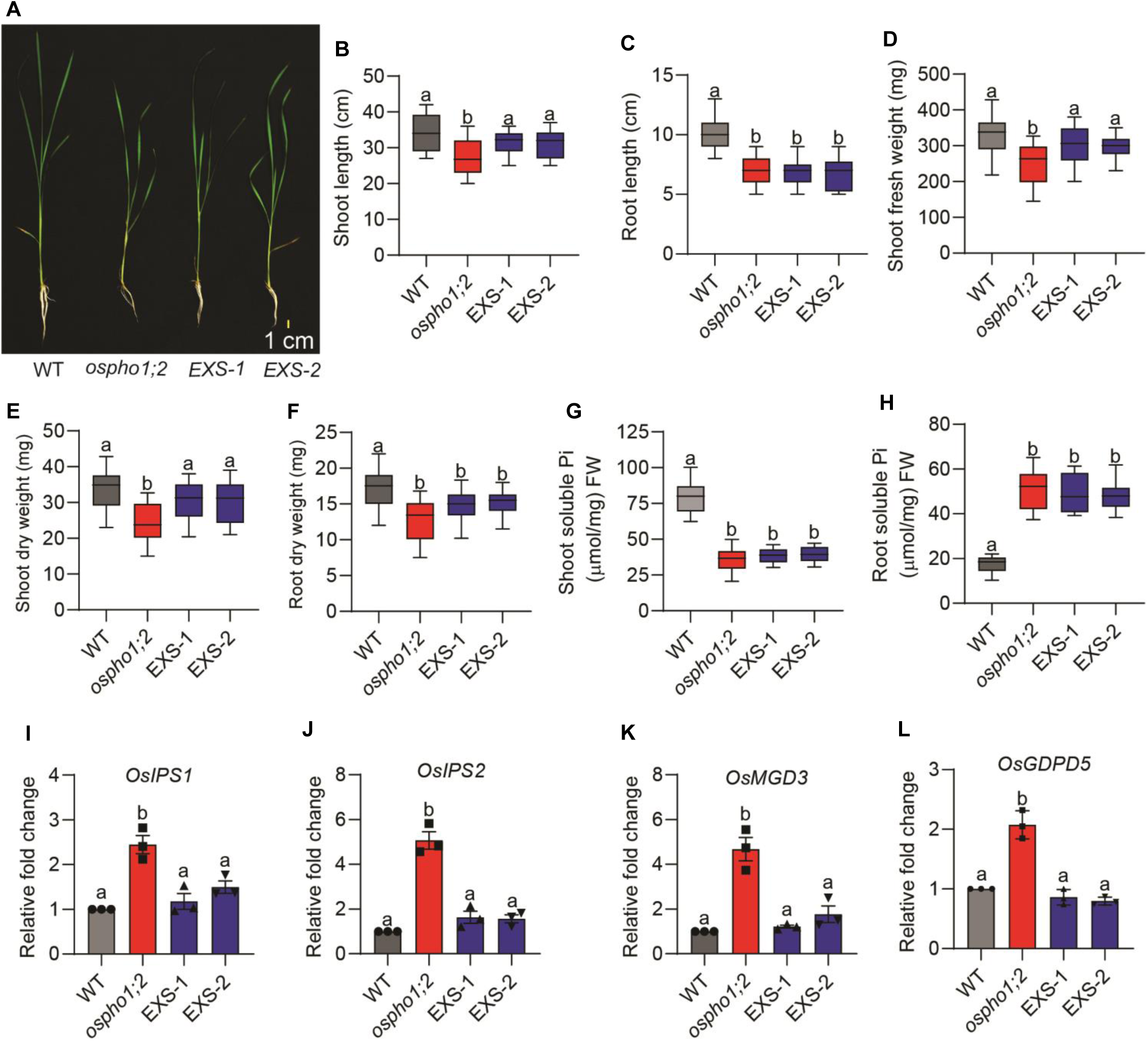
EXS lines exhibit growth similar to wild-type in Pi-sufficient conditions despite highly reduced shoot P levels. (A) The phenotypes of WT, ospho1;2, and EXS lines after 14 days of growth in Pi-sufficient (320 µm) conditions. The plant growth was estimated by measuring shoot length (B), root length (C), shoot fresh weight (D), shoot dry weight (E), and root dry weight (F). The box plot displays the mean values of 20 seedlings, with different letters on each bar indicating significant differences (ANOVA, P≤0.001). Data on soluble Pi content in the shoot (G) and root tissues (H) of WT, ospho1;2, EXS lines after 14 days of growth. The box plot shows the mean values from three biological replicates, each representing a pool of shoot tissues from 8 seedlings. Statistical differences were estimated by one-way ANOVA, with different letters on each bar indicating significant values at P≤0.05. Expression analysis of P starvation-induced (PSI) genes, including OsIPS1 (I), OsIPS2 (J), OsMGD3 (K), and OsGDPD5 (L), was analyzed in 14-day-old shoot tissues of WT, ospho1;2, and EXS lines grown for 14 days at Pi-sufficient media. The elongation factor 1-_ll_ (OsEF1_ll_) was used as an internal control for normalizing Ct values. The standard error of the mean (SEM) for three biological replicates is represented by the error bar. Statistical differences were estimated by one-way ANOVA, with different letters on each bar indicating significant values at P≤0.05.

**Fig. 3.**
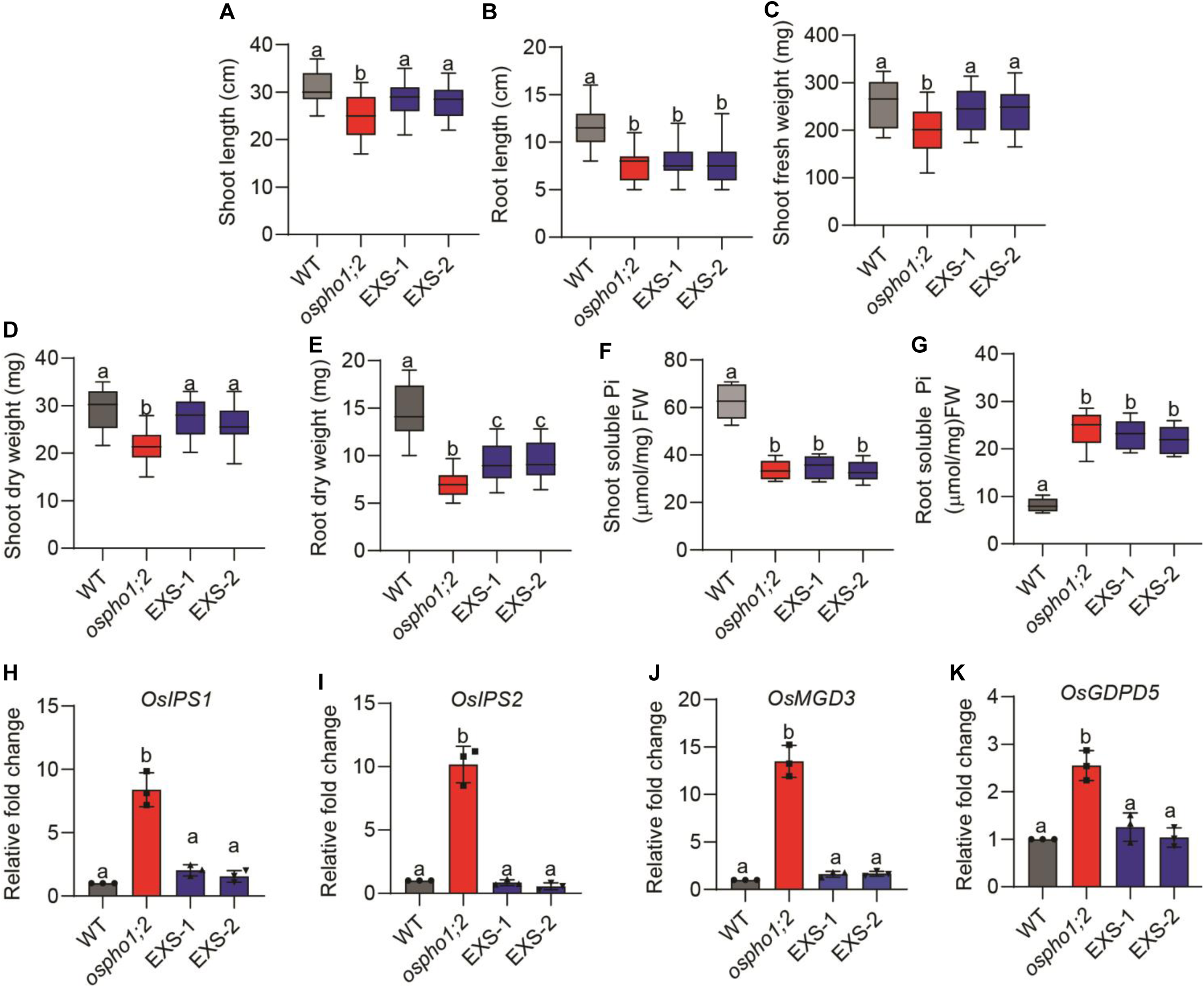
Lines expressing only the EXS domain exhibit growth similar to wild-type in Pi-deficient conditions. The WT, *ospho1;2*, and EXS line growth was estimated after 14 days in Pi-deficient (50 µM) conditions by measuring shoot length (**A**), root length (**B**), shoot fresh weight (**C**), shoot dry weight (**D**), and root dry weight (**E**). The box plot displays the mean values of 20 seedlings, with different letters on each bar indicating significant differences (ANOVA, P≤0.05). Data on soluble Pi content in the shoot (**F**) and root tissues (**G**) of WT, *ospho1;2*, EXS lines after 14 days of growth. The box plot displays the mean values of three biological replicates, each replicate representing the pool of shoot tissues from 8 seedlings. Statistical differences were estimated by one-way ANOVA, with different letters on each bar indicating significant values at P≤0.05. Expression analysis of P starvation-induced (PSI) genes, including *OsIPS1* (**H**), *OsIPS2* (**I**), *OsMGD3* (**J**), and *OsGDPD5* (**K**), was analyzed in 14-day-old shoot tissues of WT, *ospho1;2*, and EXS lines grown for 14 days on Pi-deficient media. The elongation factor 1-_ll_ (*OsEF1*_U_) was used as an internal control for normalizing Ct values. The standard error of the mean (SEM) for three biological replicates is represented by the error bar. Statistical differences were estimated by one-way ANOVA, with different letters on each bar indicating significant values at P≤0.05.

### Lines expressing 4TM+EXS (T-EXS) exhibit growth similar to *ospho1;2*

During this work, the size of the EXS domain was recently redefined (Fang et al., 2025). It now carries a few additional amino acids, including transmembrane domain 5a (TM5a), which has an amino acid, aspartic acid (D) at position 560, predicted to be essential for Pi binding and transport (**Fig. S5**). We generated the EXS line based on the previously defined size, and our lines lack TM5a. To include TM5a and find out whether this uncoupling of growth and Pi deficiency was specific to the EXS domain, we created an additional variant of *OsPHO1;2* carrying both 4TM and EXS domains, including TM5a (T-EXS). This was achieved by deleting approximately 2.2 kb region (between the first and sixth intron) of *OsPHO1; 2* that encodes the SPX domain (**Fig. S6A, B**). We generated and genotyped over 100 transgenic rice lines, ultimately identifying two lines with the desired T-EXS genotypes, named T-EXS-1 and T-EXS-2 (**Fig. S6C**). Sanger sequencing showed that the T-EXS-1 and T-EXS-2 lines had deletions of 2115 bp and 2114 bp, respectively (**Fig. S6F**). The T-EXS lines produced a truncated transcript of about 1.5 kb of *OsPHO1;2* and domain-specific expression (**Fig. S6D, E**). These lines exhibited poorer growth with shorter shoot and root lengths, as well as reduced shoot and root FW, compared to the WT. Notably, all growth parameters of the T-EXS lines were similar to those of the *ospho1;2* lines (**Fig. S7A-D**). Further, the T-EXS lines showed shoot and root Pi levels similar to those in the EXS lines and *ospho1;2* (**Fig. S7E, F**). Under Pi-deficient conditions, their growth, biomass accumulation, and Pi levels were similar to those of *ospho1;2* but significantly lower than the WT (**Fig. S8**).

Furthermore, we analyzed the growth patterns of these lines grown under Pi-sufficient conditions for 14 days (**Fig. 4**). The T-EXS lines continued to show growth, biomass, and Pi levels similar to those of *ospho1;2* (**Fig. 4A-H**). Interestingly, RT-qPCR analysis revealed that PSI genes, such as *OsIPS1*, *OsIPS2*, *OsMGD3*, and *OsGDPD5*, were not upregulated in the shoot tissues of T-EXS compared to WT (**Fig. 4I-L**). These lines displayed the same trend when grown in Pi-deficient conditions (**Fig. S9**). These data collectively indicate that the expression of the T-EXS domain, which includes TM5a, does not facilitate Pi transport in shoot tissues and fails to produce the growth improvements observed in the EXS lines.

**Fig. 4.**
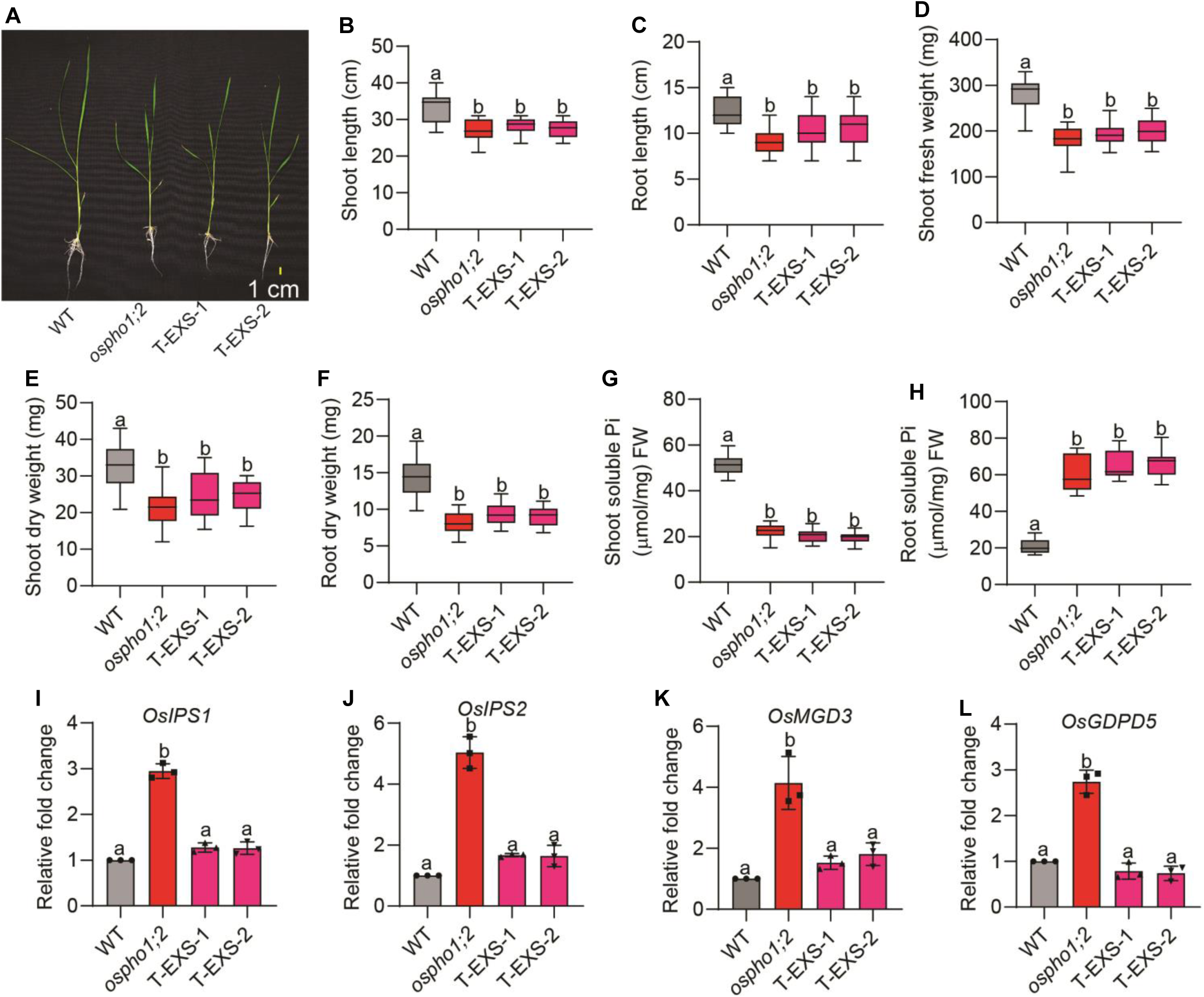
Lines expressing the 4TM+EXS domain (T-EXS) behave like *ospho1;2* mutants under Pi-sufficient conditions. **(A)** The phenotypes of WT, *ospho1;2*, and T-EXS lines after 14 days of growth in Pi-sufficient (320 µm) conditions. The plant growth was estimated by measuring shoot length (**B**), root length (**C**), shoot fresh weight (**D**), shoot dry weight (**E**), and root dry weight (**F**). The box plot displays the mean values of 20 seedlings, with different letters on each bar indicating significant differences (ANOVA, P≤0.05). Data on soluble Pi content in shoot tissues (**G**) and root tissues (**H**) of WT, *ospho1;2*, and T-EXS lines after 14 days of growth. The box plot displays the mean values of three biological replicates, each representing a pool of shoot tissues from eight seedlings. Statistical differences were estimated by one-way ANOVA, with different letters on each bar indicating significant values at P≤0.05. Expression analysis of P starvation-induced (PSI) genes, including *OsIPS1* (**I**), *OsIPS2* (**J**), *OsMGD3* (**K**), and *OsGDPD5* (**L**), was analyzed in 14-d-old shoot tissues of WT, *ospho1;2*, and 4TM+EXS lines grown for 14 days at Pi-sufficient media. The elongation factor 1-_ll_ (*OsEF1*_U_) was used as an internal control for normalizing Ct values. The standard error of the mean (SEM) for three biological replicates is represented by the error bars. Statistical differences were estimated by one-way ANOVA, with different letters on each bar indicating significant values at P≤0.05.

### RNA-Seq analysis of WT, *ospho1;2,* and EXS lines reveals suppression of defense pathways in EXS lines

EXS lines displayed healthy shoot growth despite having a very low Pi, suggesting these lines might have altered Pi sensing/signaling. To understand the molecular mechanisms behind good growth in EXS lines, we performed RNA sequencing (RNA-Seq) on the shoots of WT, *ospho1;2*, and EXS-1 lines grown for 14 days under Pi-sufficient conditions. It is expected that both *ospho1;2* and EXS lines with Pi-deficient shoots should activate PSR. Principal component analysis (PCA) distinctly separated the WT, *ospho1;2*, and EXS lines, indicating different transcriptional responses (**Fig. S10A**). Differential gene expression analysis revealed that the *ospho1;2* mutant had significantly more differentially expressed genes (DEGs) than the EXS line. Of the 2200 DEGs identified in *ospho1;2* compared to WT, 241 were upregulated and 297 downregulated. In contrast, of the 1832 DEGS, the EXS lines showed only 116 upregulated and 549 downregulated (**Fig. 5A, Table S2**). Further, both lines had unique sets of upregulated genes, with only 9% overlapping; however, 36% of the downregulated genes were common to both *ospho1;2* and EXS lines (**Fig. S10B**). Heat map analysis showed that the *ospho1;2* mutant significantly upregulated PSI genes, such as *OsIPS1*, *OsSPX-MFS2*, and *OsGDPDs* (**Fig. 5B**). Furthermore, genes involved in jasmonic acid (JA) biosynthesis, like lipoxygenase 2.3 (*OsLOX 2.3*) and jasmonyl methyl transferase (*OsJMT*), which convert JA to methyl jasmonate (MeJA), were markedly upregulated in the *ospho1;2* mutant (**Fig. 5B**). Genes related to abscisic acid (ABA) synthesis, including *OsNCED1*, *OsNCED2*, and the ABA receptor PYL10-like, also showed significant upregulation, indicating elevated hormonal signaling in response to Pi deficiency (**Fig. 5B**). Conversely, the EXS lines showed no upregulation of the PSI genes or those involved in JA or ABA biosynthesis. The EXS lines showed increased expression of several transcription factors, such as *OsMADS61* and *OsHSFC1a*, along with proteins like lipid transfer protein (*OsLTP47*), burp-domain containing protein (*OsBURP8*), LRR-receptor-like kinase (OsRLCK258), and JA-upregulated protein 1 (*OsJAUP1*) (**Fig. 5C**). Many downregulated genes in *ospho1;2* were linked to defense responses, including terpene synthases (*OsTPS46, OsTPS30, OsKSL4*), pathogenesis-related genes (*OsPR10a, OsPR10c*), WRKY transcription factors (*OsWRKY65, OsWRKY89*), and cytochrome enzymes (*OsCYP76M10, OsCYP51H5*) (**Fig. S11A**). Compared to *ospho1;2*, the EXS lines exhibited greater suppression of terpene synthases (*OsTPS3, OsTPS46, OsTPS30, OsCPS4*), WRKY domain proteins (*OsWRKY40, OsWRKY62, OsWRKY65, OsWRKY89*), and pathogenesis-related genes (*OsPR5, OsPR10a, OsPR10c*) (**Fig. S11B**). This suggests that reducing defense responses allows more resources for plants to face Pi deficiency in EXS lines. Gene Ontology (GO) and KEGG pathway analyses revealed different biological processes affected in *ospho1;2* and EXS lines (**Fig. 5D, E**). In *ospho1;2*, upregulated genes were involved in carotenoid dioxygenase activity, 9-cis-epoxycarotenoid dioxygenase activity, transmembrane transport, and iron-ion transport. The upregulated genes in EXS lines did not show significant enrichment in known GO terms. KEGG analysis identified that genes related to photosynthesis, carotenoid biosynthesis, and linoleic acid metabolism were enriched in *ospho1;2*. Likewise, genes involved in alcohol biosynthesis, small molecule biosynthesis, and secondary metabolites were enriched in EXS lines (**Fig. 5D**). Downregulated genes in both lines were associated with stress response, defense response, diterpenoid synthesis, terpene synthase activity, carbohydrate binding, and receptor signaling pathways, with these effects being more pronounced in the EXS lines. KEGG pathway analysis showed enrichment in plant-pathogen interactions and phenylpropanoid biosynthesis specifically in *ospho1;2*, while alpha-linolenic acid metabolism was enriched in EXS lines (**Fig. 5E**). Unlike the upregulated genes, many GO terms and KEGG pathways among the downregulated genes were shared by both lines. These results imply that suppression of PSI and defense-related hormone genes underpins the enhanced growth observed in EXS lines compared to *ospho1;2*.

**Fig. 5.**
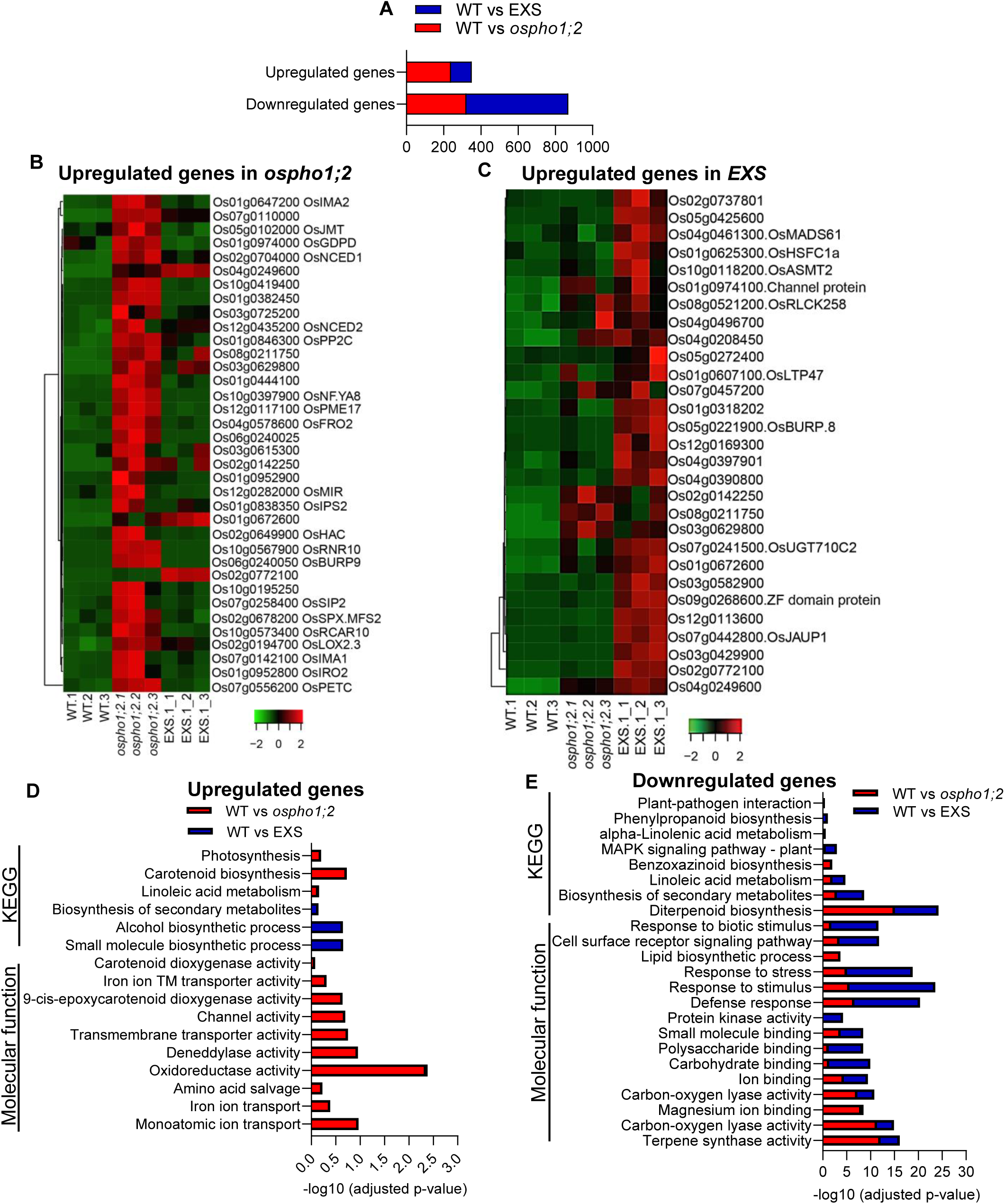
RNA-Seq analysis of *ospho1;2* genome-edited lines. The WT, *ospho1;2*, and EXS (expressing the EXS domain of *OsPHO1;2*), line seeds were germinated in ½ MS media for 5 days and then transferred to Yoshida media containing 320 µM phosphate (Pi sufficient) for 14 days, and shoot tissue was harvested for RNA-Seq analysis. (**A**) The total number of significantly expressed genes (DEGs) (Log2FC-1/+1, FDR 0.05) in *ospho1;2* and EXS line compared to WT. Heat map representing the normalized expression values of the top 25 upregulated genes, PSI and ABA biosynthetic genes in *ospho1;2* (**B**), and the top 30 upregulated genes in EXS (**C**) lines. Top 20 significantly enriched Gene Ontology (GO) terms and KEGG pathways of upregulated DEGs (**D**) and downregulated DEGs (**E**) in *ospho1;2* and EXS lines. GO terms were defined as significantly enriched if the false discovery rate (FDR) was ≤0.05.

### Phytohormones profiling suggests a role for JA and ABA in EXS lines

RNA-seq data suggested that Pi deficiency significantly affects hormonal signaling (JA, ABA) pathways in *ospho1;2* mutants. The phytohormone profiling was done in the shoot of WT, *ospho1;2*, and EXS lines grown under Pi-sufficient conditions. Jasmonyl isoleucine (JA-Ile), a bioactive form of JA, showed a substantial increase in *ospho1;2* mutants compared to WT (**Fig. 6A**). Interestingly, JA-Ile levels in the EXS line were significantly lower than in *ospho1;2* but similar to those in WT (**Fig. 6B**). Likewise, ABA levels were higher in the *ospho1;2* mutant than in both WT and EXS lines (**Fig. 6B**). This revealed that expressing the EXS domain is sufficient to prevent excess JA/ABA levels despite the low shoot Pi levels. Further, the expression of JA biosynthetic genes, including *OsLOX1*, *OsLOX2*, *OsAOC*, and *OsAOS1*, was significantly increased in *ospho1;2* mutants but remained at the WT levels in EXS lines (**Fig. 6C-G**). Similarly, the expression of ABA biosynthetic genes, *OsNCED1*, *OsNCED2*, and *OsNCED3*, was significantly higher in the *ospho1;2* mutants compared to WT and EXS lines (**Fig. 6H-K**). Although the expression of PSI genes was suppressed, the growth of T-EXS lines was similar to that of *ospho1;2*. To determine if hormone signaling was affected in T-EXS lines, we measured the levels of JA-Ile and ABA in their shoots. Both JA-Ile and ABA contents were nearly twice as high in T-EXS lines compared to WT, but similar to *ospho1;2* lines (**Fig. 6A, B**). Additionally, the expression levels of JA and ABA biosynthetic genes were significantly increased in T-EXS lines relative to WT (**Fig. 6C-K**). These results suggest that Pi deficiency caused by the loss of *ospho1;2* activates defense hormone pathways, especially JA-Ile and ABA. The ability of the EXS domain to restore hormone levels and gene expression to WT levels may support improved plant growth. In contrast, the 4TM+EXS domains failed to reduce these hormone levels in Pi-deficient shoots, which negatively affects plant growth.

**Fig. 6.**
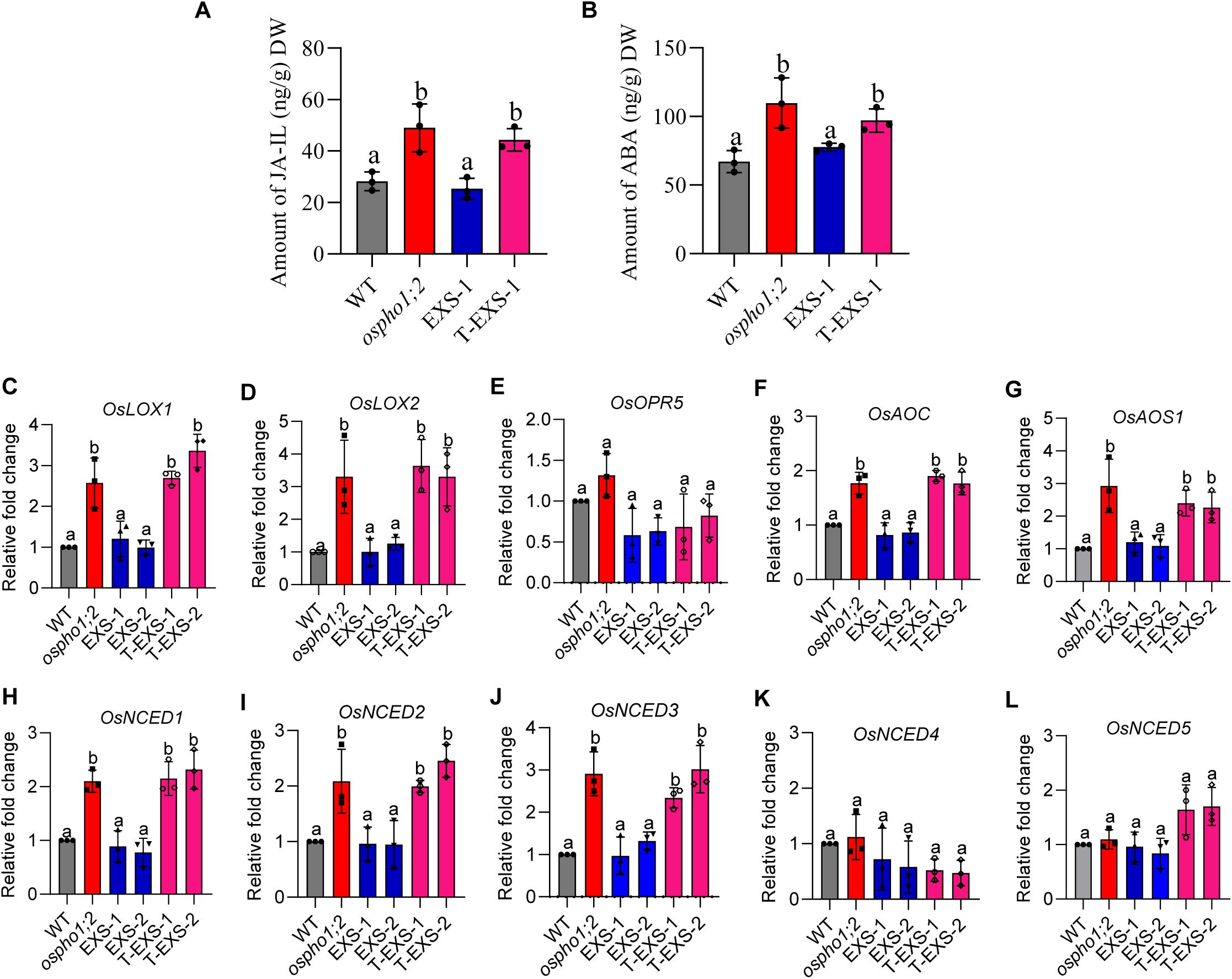
Phytohormone profiling indicates a role for JA and ABA in the EXS lines. (**A**) Jasmonyl isoleucine (JA-Ile) and (**B**) abscisic acid (ABA) content in the shoot tissues of WT, *ospho1;2*, EXS, and T-EXS lines after 14 days of growth in Pi-sufficient conditions (320 µM Pi). The data represent the average of 3 independent replicates, each with at least 10 seedlings. One-way ANOVA was performed to estimate the statistical difference among the different groups. Different letters at each bar indicate the statistical significance at P value ≤0.05. (**C-G**) Expression levels of jasmonic acid (JA) biosynthetic genes, including *OsLOX1*, *OsLOX2*, *OsOPR5*, *OsAOC*, and *OsAOS1* in shoot tissues. The elongation factor 1-_ll_ (*OsEF1*_U_) was used as an internal control for normalizing Ct values. The standard error of the mean (SEM) values for three biological replicates are represented by the error bars, with different letters on each bar indicating significant differences (ANOVA, P≤0.05). (**H-L**) Expression levels of ABA biosynthetic genes, including *OsNCED1*, *OsNCED2*, *OsNCED3*, *OsNCED4*, and *OsNCED5* in shoot tissues. The elongation factor 1-_ll_ (*OsEF1*_U_) was used as an internal control for normalizing Ct values. The standard error of the mean (SEM) values for three biological replicates are represented by the error bars, with different letters on each bar indicating significant differences (ANOVA, P≤0.05).

### EXS domain partially rescues the phenotype of *ospho1;2* at the mature stage

The phenotypes of *ospho1;2*, EXS, and T-EXS were examined in mature plants. The *ospho1;2* line shows significant reductions in key agronomic traits such as plant height, tiller number, leaf width, and flag leaf width (**Fig. S12A-F**). Unlike at the seedling stage, the EXS lines did not maintain growth comparable to WT, while the T-EXS lines continued to behave like *ospho1;2*. However, the EXS lines show improved plant height, tiller number, leaf width, and flag leaf width when compared to *ospho1;2* (**Fig. S12A-F**). Functional OsPHO1;2 is needed for proper seed development in rice (Ma *et al.,* 2021; Maurya *et al.,* 2025). Moreover, *ospho1;2*, EXS, and T-EXS lines displayed visibly shrunken and poorly developed seeds compared to WT (**Fig. 7A**). A significant decrease in seed width and 100-seed weight was observed in these lines compared to WT. Notably, the 100-seed weight in the EXS and T-EXS lines is significantly higher than in *ospho1;2* (**Fig. 7B-D)**. The total P content in the caryopsis of WT is considerably higher than in *ospho1;2*, EXS, and T-EXS lines (**Fig. 7E**). Similarly, the total P content of brown WT seeds, containing ∼20 mmol of P, is higher compared to only ∼10 mmol in the *ospho1;2,* EXS, and T-EXS seeds (**Fig. 7F**). Interestingly, the opposite trend was observed in the husk, where *ospho1;2,* EXS, and T-EXS lines have significantly higher total P compared to WT (**Fig. 7G**). However, seeds from all these lines germinate well like WT, indicating that seed defects in these genotypes didn’t impair seed germination (**Fig. S13**). Collectively, these data suggest that full-length *OsPHO1;2* is crucial for maintaining plant growth at the mature stage and seed development. The expression of the EXS domain alone failed to support plant growth during later stages or seed development.

**Fig. 7.**
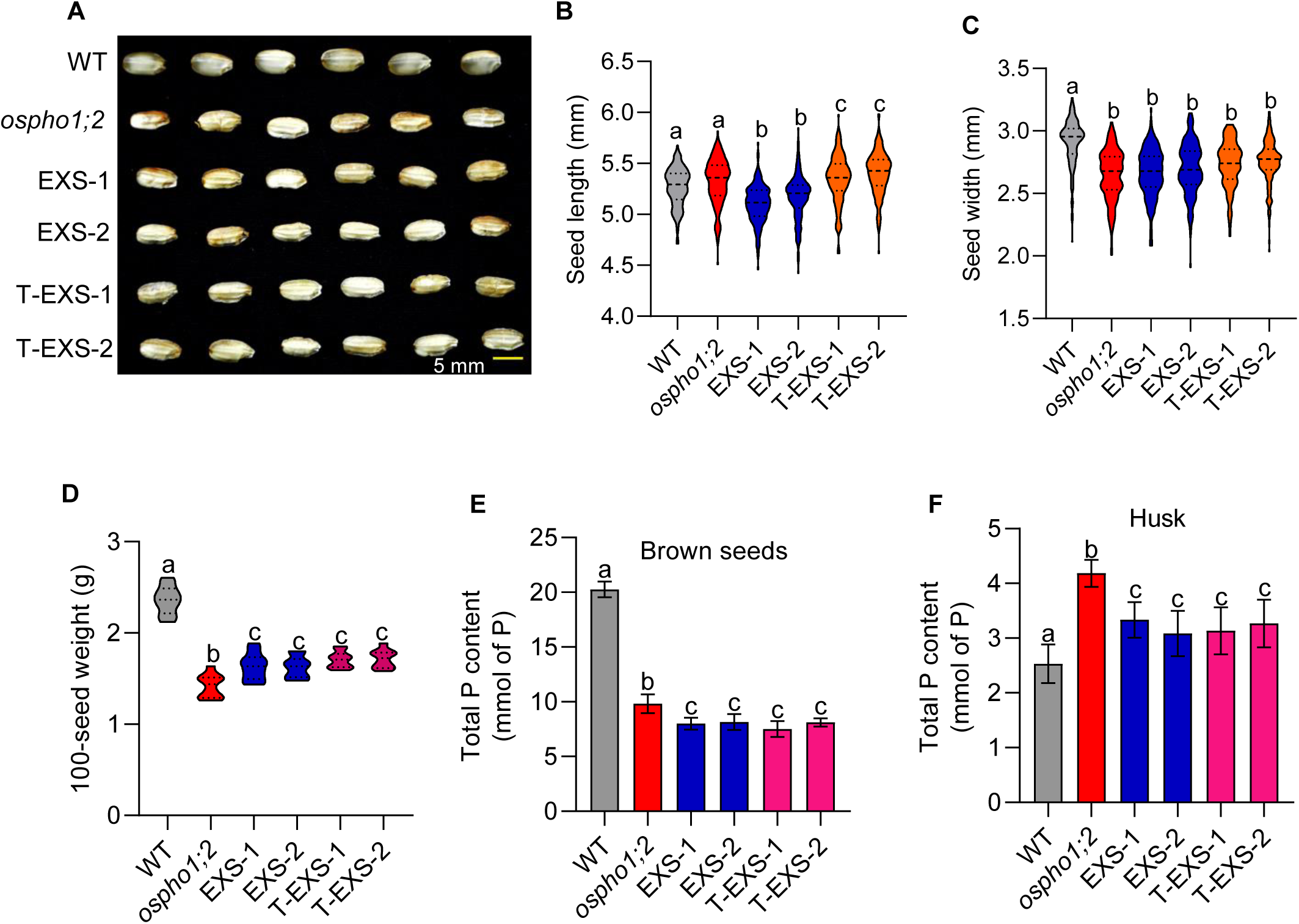
Seed phenotypic analysis of *OsPHO1;2* genome-edited lines. (**A**) Mature dehusked seed morphology of WT, *ospho1;2*, EXS, and T-EXS lines. The violin plot represents the seed length (**B**), seed width (**C**), and 100-seed weight (**D**) of *OsPHO1;2* genome-edited lines. The data display the mean values from at least 100 seeds, with different letters on each bar indicating significant differences (ANOVA, P ≤ 0.05). Data on total P content in the caryopsis (**E**), brown seeds (**F**), and husk (**G**) of WT, *ospho1;2*, EXS, and T-EXS lines. Statistical differences were estimated by one-way ANOVA, with different letters on each bar indicating significant values at P≤0.05 (n=3).

### A single allele of *OsPHO1;2* is sufficient for maintaining plant growth and seed development

EXS lines grow well while suppressing PSR despite low shoot Pi levels; however, the poor shoot Pi hampers yield. We hypothesize that having one functional allele of *OsPHO1* in a heterozygous plant may restore the yield. Therefore, we decided to examine the behavior of heterozygous plants of OsPHO1;2/*ospho1;2* (Pp) lines in terms of plant growth, Pi transport, and seed development. Since the T-EXS lines mimic the growth behavior of the *ospho1;2* mutant, heterozygous plants of these lines were used for the study (**Fig. 8A**). The Pp lines displayed shoot length, root length, seedling fresh weight, seedling dry weight, and shoot and root Pi levels comparable to WT, but better than those of *ospho1;2* (**Fig. 8B-H**). This indicates that a single PHO1 allele is enough to maintain plant growth and Pi transport. Additionally, seed morphology analysis revealed that Pp lines produced healthy seeds like WT (**Fig. 8I, J**). Moreover, the total P content in the brown seeds and husk of these lines was also similar to WT (**Fig. 8K, L**). Overall, these findings suggest that one functional PHO1 allele is sufficient for Pi export from roots to shoots and from maternal to filial tissues in developing seeds, thereby restoring normal plant growth and seed development.

**Fig. 8.**
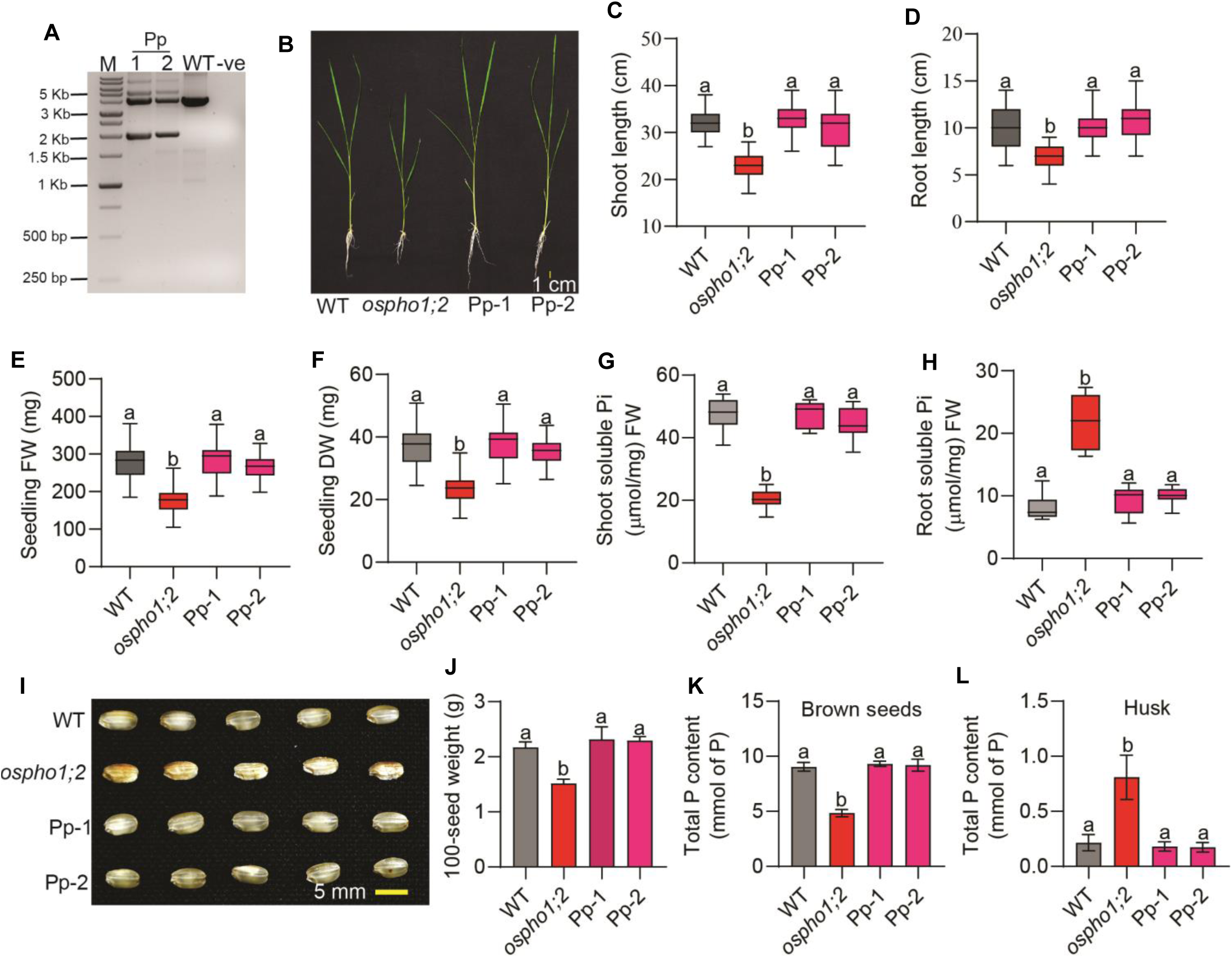
A single allele of *OsPHO1;2* is sufficient for maintaining plant growth and seed development. **(A)** Genotyping of progenies to select Pp (OsPHO1;2/T-EXS) by genomic DNA-PCR. M-DNA ladder; bp-base pairs,-ve - PCR negative. (**B**) The phenotypes of WT, *ospho1;2*, and Pp lines after 14 days of growth in Pi-sufficient (320 μM) conditions. The plant growth was estimated by measuring shoot length (**C**), root length (**D**), seedling fresh weight (**E**), and seedling dry weight (**F**). The box plot displays the mean values of 20 seedlings, with different letters on each bar indicating significant differences (ANOVA, P≤0.001). Data on soluble Pi content in the shoot (**G**) and root tissues (**H**) of WT, *ospho1;2*, and Pp lines after 14 days of growth. The box plot displays the mean values of three biological replicates, each representing a pool of shoot tissues from eight seedlings. Statistical differences were estimated using one-way ANOVA, with different letters on each bar indicating significant differences at P ≤ 0.05. (**I**) Mature dehusked seed morphology of WT, *ospho1;2*, and Pp lines. (**J**) The 100-seed weight of *OsPHO1;2* genome-edited lines. The data display the mean values from at least 100 seeds, with different letters on each bar indicating significant differences (ANOVA, P ≤ 0.05). Data on total P content in the brown seeds (**K**), and husk (**L**) of WT, *ospho1;2*, and Pp lines. Statistical differences were estimated using one-way ANOVA, with different letters on each bar indicating significant values at P ≤ 0.05 (n = 3).

## Discussion

Plants accumulate P at varying levels across different crops, with concentrations typically ranging from 0.05% to 0.5% of their dry weight (Wieczorek et al., 2022). Crops like rice, corn, and wheat, which require medium P (15 mg kg ¹ Olsen P), can accumulate 0.2% to 0.5% of their dry weight at various growth stages (Wang et al., 2023; McDowell et al., 2024). Interestingly, rice seeds with low P content can grow well without any growth defects (Rose et al., 2012; Rose & Raymond, 2020; Yugandhar et al., 2022). Genome-editing of the SULTR-like phosphorus distribution transporter (SPDT), which regulates P allocation to the grain, results in a 20-30% reduction in seed P without affecting yield, seed germination, or seedling vigor (Yamaji et al., 2017). These finding suggests that even under low seed P content, plants can grow well without Pi-deficiency symptoms. In this study, we demonstrate that growth retardation caused by low Pi can be separated by expressing only the EXS domain of OsPHO1;2.

All three lines of *ospho1;2*, EXS and T-EXS displayed a significantly reduced shoot Pi levels. Recent structural work using cryo-EM on AtPHO1;H1, a functional paralogue of AtPHO1 (Stefanovic et al., 2011), as well as of the human orthologue XPR1, have shown that PHO1/XPR1 form a homodimeric Pi channel (Yan et al., 2024; Lu et al., 2024; Fang et al., 2025; Chen et al., 2025; Zhang et al., 2025). The N-terminal SPX domain binding IP7/IP8 is followed by a series of transmembrane helices, which terminate in a hydrophilic cytosolic C-terminal domain. These structural studies show that the first 4 transmembrane helices (4TM) are important for dimerization, while the EXS domain forms the Pi pore containing six TM helices. The binding of IP7/IP8 to the SPX domain leads to stabilization of the dimeric structure and opening of the pore. Thus, removal of the SPX domain from either PHO1 or XPR1 leads to unstable non-functional proteins when expressed in their endogenous tissues, although overexpression of a AtPHO1 TM-EXS constructs in heterologous tissues may still show some Pi transport activity (Wege et al., 2016). In agreement with the requirement of a functional SPX domain, point mutants in the SPX domain of AtPHO1, which abolish IP7/IP8 binding, fail to complement the low Pi phenotype of the *atpho1* mutant (Wild et al., 2016). Expression of only the EXS domain of AtPHO1 also failed to show any Pi export activity (Wege et al., 2016). Thus, the effects of the expression of either the EXS domain only or of the 4TM-EXS domain of OsPHO1;2 are unlikely to be explained by changes in Pi transport activity, thus suggesting a signaling role for the EXS domain.

Plants with low Pi levels activate phosphate starvation response (PSR), which leads to the activation of PSI genes and thereby affects morphological, physiological, and molecular processes (Wang et al., 2014; Bouain et al., 2016). The *atpho1* mutants are shown to activate PSI genes in the shoot even under Pi-sufficient conditions due to defects in Pi transport from root to shoot (Wang et al., 2004; Secco et al., 2010; Rouached et al., 2011). RNA-Seq and RT-qPCR revealed that a significant set of genes associated with PSR showed decreased expression in the EXS lines. Furthermore, RNA-Seq data revealed that most of the downregulated genes in EXS lines are associated with the defense response. Under Pi deficiency, plants adopt sophisticated resource allocation strategies that favor growth maintenance over defense capabilities. This adaptive strategy represents a key trade-off where plants strategically lower defense response to maximize growth under Pi-limited conditions (Huot et al., 2014; Karasov et al., 2017; Inoue et al., 2024; Zrimec et al., 2025). This indicates that the EXS domain can suppress or modulate the systemic signaling that typically activates these responses under Pi starvation, thereby maintaining better plant growth.

Plants with low Pi levels accumulate higher levels of defense phytohormones such as JA and ABA (Khan et al., 2016; Zhang et al., 2022b; Lei et al., 2022; Jaskolowski & Poirier, 2024). The persistent activation of the JA response is growth-suppressing (Yang et al., 2012; Huot et al., 2014; Jin et al., 2023). Similarly, increased levels of ABA inhibit plant growth and productivity but improve tolerance to various environmental stresses (Cutler et al., 2010; Kavi Kishor et al., 2022). The _а_*tph*_о_*1* mutant grown under Pi-sufficient conditions was shown to accumulate higher amounts of JA and Jasmonyl isoleucine (JA-Ile) (Khan et al., 2016; Jaskolowski & Poirier, 2024). This work shows a close correlation between levels of JA and ABA in Pi-deficient plants and growth. While the EXS lines are Pi-deficient, they show WT-like growth at the early stages and low JA and ABA levels. In contrast, the T-EXS lines show equally low Pi content but have higher ABA and JA levels and show reduced growth similar to the *ospho1;2* mutant. These data suggest that suppressed defense responses might help the EXS lines perform better at the early vegetative stage. It would be important to further investigate how the presence of the EXS domain, alone or in combination with the 4TM-EXS domain, can suppress the defense response in rice. In Arabidopsis, both 4TM-EXS and EXS are localized to the Golgi/trans-Golgi (Wege et al., 2016). It is possible that the expression of EXS alone leads to modifications in the interaction of proteins at the endosomal and plasma membranes that are important for JA and ABA signaling.

The EXS lines behave like *ospho1;2* mutants in later development stages. Additionally, both lines produced shrunken, poorly developed seeds similar to those of *ospho1;2*. This mainly results from defective Pi mobilization during grain filling, as indicated by significantly higher Pi levels in the husk (Ko *et al*., 2024). In Arabidopsis, *PHO1* is implicated in transferring Pi from the chalazal seed coat to the embryo (Vogiatzaki *et al*., 2017). Similarly, in crops like rice and maize, *pho1;2* mutants showed defects in seed development and yield due to impaired Pi export from maternal to filial tissues (Ma *et al*., 2021; Ko *et al*., 2024). The link between lower seed P content and reduced seed size and weight confirms that *OsPHO1;2*-mediated Pi transport is vital for seed development and grain filling in rice. This stage-specific limitation suggests that different developmental phases may require distinct PHO1 domain functions or that prolonged Pi deficiency may eventually outweigh the growth benefits of EXS domain expression. Furthermore, this indicates that reducing PSR signaling or disconnecting growth defects from Pi deficiency symptoms alone isn’t enough to sustain plant performance regarding yield. This highlights the essential physiological role of phosphate in plant growth and development. Interestingly, Pp lines developed healthy seeds like WT. This suggests that developing hybrid lines with one functional PHO1 allele offers an alternative approach to fine-tune, rather than fully disrupt, transporter activity.

P fertilizers are crucial for rice production, requiring at least 15 mg kg ¹ of Olsen P for optimal growth. P is a non-renewable resource from limited rock phosphate reserves, with current global reserves potentially lasting 200-400 years, depending on demand scenarios. However, phosphorus use efficiency (PUE) is low, with rice utilizing only about 25% of applied fertilizers, necessitating excessive P applications in Pi-deficient soils (Lynch, 2011; Haefele et al., 2014). Rice PUE can be enhanced by engineering Pi transporters, modulating transcriptional and post-translational regulation, strengthening the internal P use efficiency, and establishing a symbiotic relationship with beneficial bacteria (Heuer et al., 2017; Paz-Ares et al., 2022; Guan et al., 2022; Lu et al., 2023; Zulfiqar et al., 2025). In this study, we propose that the expression of only the EXS domain of *OsPHO1;2* reduces the activity of PSR signaling and the activation of defense phytohormones, which leads to enhanced plant growth. This finding opens new avenues for understanding the mechanistic role of the EXS domain in mediating Pi signaling. Furthermore, this research enhances our understanding of how different domains of *OsPHO1;2* work at the molecular level and supports the development of genetic strategies to increase crop phosphate efficiency, a vital factor for sustainable agriculture in phosphate-deficient environments.

## Author contributions

B.M. conducted experiments, analyzed data, and wrote the first draft. K.M., LV, PG, PSK, GG, and AJ helped with the experiments. J.G. and Y.P. conceived the project; J.G. supervised the experiments, analyzed the data, and finalized the manuscript with assistance from Y.P.

## Acknowledgments

This research is funded by the Swarnajayanti fellowship (SB/SJF/2019-20/07) and Indo-Swiss research grant (BT/IN/Swiss/46/JG/2018-2019) to JG and a grant from the Herbette Foundation at the University of Lausanne to YP. Research fellowships to B.M. from DBT, India, K.M. from CSIR, India, and U.S. from NIPGR, India, are gratefully acknowledged.

## Data availability statement

All relevant data generated or analysed are included in the manuscript with supporting materials.

## Figure Legends

**Fig. S1. SDS-PAGE analysis of purified SpCas9-His protein.**

**Fig. S2. *ospho1;2* CRISPR/Cas9 mutants are deficient in Pi export.**

**Fig. S3. Rice lines expressing the EXS domain grew like WT under Pi-sufficient conditions.**

**Fig. S4. Rice lines expressing the EXS domain grew like WT under Pi-deficient conditions.**

**Fig. S5. Diagrammatic representation of OsPHO1;2 protein.**

**Fig. S6. CRISPR/Cas9-mediated genome-editing of *OsPHO1;2* to express 4TM+EXS domain under the *OsPHO1;2* native promoter (T-EXS lines).**

**Fig. S7. Lines expressing the 4TM+EXS domain (T-EXS) grew like *ospho1;2* under Pi-sufficient conditions.**

**Fig. S8. Lines expressing the 4TM+EXS domain (T-EXS) grew like *ospho1;2* under Pi-deficient conditions.**

**Fig. S9. Lines expressing the 4TM+EXS (T-EXS) domain behave like *ospho1;2* mutants under Pi-deficient conditions.**

**Fig. S10. RNA-Seq analysis of *ospho1;2* genome-edited lines.**

**Fig. S11. Heat map analysis of downregulated genes of *ospho1;2* genome-edited lines.**

**Fig. S12. Phenotypic analysis of *OsPHO1;2* genome-edited lines at the mature stage.**

**Fig. S13. *OsPHO1;2* genome-edited lines germinate like WT.**

